# Macrophage transplantation rescues RNASET2-deficient leukodystrophy by replacing deficient microglia in a zebrafish model

**DOI:** 10.1101/2023.12.04.569924

**Authors:** Holly A Rutherford, Diogo Candeias, Christopher JA Duncan, Stephen A Renshaw, Noémie Hamilton

**Author notes:** Corresponding author: Noémie Hamilton, York Biomedical Research Institute Department of Biology, University of York Wentworth Way York, YO10 5DD, United Kingdom.

## Abstract

RNaseT2-deficient leukodystrophy is a rare infantile white matter disorder mimicking a viral infection and resulting in severe psychomotor impairments. Despite its severity, there remain no treatments for this disorder, with little understanding of cellular mechanisms of pathogenesis. Recent research using the *rnaset2* mutant zebrafish model has suggested that microglia – brain-resident phagocytes – may be the drivers of neuroinflammation in this disorder, due to their failure to digest apoptotic debris during early development. As such, the current study aimed to develop a strategy for microglial replacement and test the effects of this intervention on *rnaset2* mutant zebrafish pathology. We developed a strategy for microglial replacement through transplantation of adult whole kidney marrow-derived macrophages into embryonic hosts. Using live imaging, we revealed that transplant-derived macrophages can engraft within host brains and express microglia-specific markers, suggesting adoption of a microglial phenotype. Tissue clearing strategies revealed the persistence of transplanted cells in host brains beyond embryonic stages We demonstrated that transplanted cells clear apoptotic cells within the brain, as well as rescuing overactivation of the antiviral response otherwise seen in mutant larvae. RNA sequencing at the point of peak transplant-derived cell engraftment confirms that transplantation can reduce the brain-wide immune response, and particularly the antiviral response, in *rnaset2*-deficient brains. Crucially, this reduction in neuroinflammation resulted in behavioural rescue - restoring *rnaset2* mutant motor activity to wild type levels in embryonic and juvenile stages. Together, these findings demonstrate the role of microglia as the cellular drivers of neuropathology in *rnaset2* mutants, and that macrophage transplantation is a viable strategy for microglial replacement in the zebrafish. Therefore, microglia-targeted interventions may have therapeutic benefits in RNaseT2-deficient leukodystrophy.

## Introduction

As the resident immune cells of the brain, microglia are essential to the development and function of a healthy nervous system. These highly specialised tissue-resident phagocytes have a host of functions involved in CNS homeostasis, including phagocytosing debris and dying cells, mediating the immune response through release of carefully balanced pro- and anti-inflammatory mediators, and supporting the health of neurons and other glial cells [7, 17, 24, 26, 36, 53]. It is perhaps unsurprising, therefore, that microglial dysfunction has been linked to a multitude of disorders, including Alzheimer’s disease, Parkinson’s disease and white matter diseases [14, 27, 29, 37, 39]. Although white matter is composed predominantly of myelin - an insulating protein produced and stored within the expansive membranes of oligodendrocytes - it is becoming increasingly apparent that microglial dysfunction can contribute to such diseases, both by directly impacting myelin health and indirectly, by promoting neuroinflammation.

One family of white matter diseases in which the contribution of microglia is becoming more apparent are the leukodystrophies - a family of monogenic white matter disorders, many of which present during infancy [27, 46]. These devastating conditions are associated with extensive psychomotor impairment, the formation of white matter lesions, neuroinflammation and, in many cases, limited life expectancy. Although the mechanisms of pathology are, in many instances, specific to the gene affected in each leukodystrophy, microglial dysfunction has begun to emerge as a common theme - even in disorders in which the mutation is thought to affect a gene primarily associated with the function of other glial cell types [2, 27, 40, 49]. In particular, microglial activation has been suggested to be the first marker - and therefore potential driver - of pathology in animal models of multiple leukodystrophies, which has been corroborated by several post-mortem reports from patients [2, 38, 51]. In addition, haematopoietic stem cell (HSC) transplantation – which may act to replace microglia – is a clinically established therapy in several leukodystrophies [3, 5, 12, 13, 18, 34, 42, 48]. As such, microglia-targeted approaches may represent a promising therapeutic avenue in the leukodystrophies.

Here, we show that the addition of healthy microglia reduces both neuropathology and symptom presentation in a zebrafish model of RNASET2-deficient leukodystrophy – a disorder which sits at the intersection of leukodystrophies, lysosomal storage diseases and a family of autoinflammatory disorders characterised by upregulation of the interferon response, known as interferonopathies [21, 23, 25]. We have previously demonstrated that a failure of *rnaset2* mutant microglia to clear developmental apoptosis contributes to downstream pathology - leading to an elevated antiviral response, behavioural abnormalities and reduced survival in the zebrafish model [21]. We therefore hypothesize that microglia are the cellular drivers of neuropathology in this disorder, and that replacing these deficient cells will rescue *rnaset2* mutant phenotypes. In this study, we develop a methodology to replace deficient microglia by macrophage transplantation and demonstrate that transplantation of macrophages can replenish the microglial population in both healthy and *rnaset2* mutant zebrafish. Moreover, replacing deficient microglia in *rnaset2* mutants is sufficient to rescue cell death, antiviral immune responses and defects in locomotion during embryonic stages. RNA sequencing and behavioural assays confirmed the therapeutic effect of transplanted cells in later stages, with complete rescue of neuroinflammation and impaired swimming. As such, our findings provide further evidence that microglia are central to RNASET2-deficient leukodystrophy pathology and support a targeting of microglia as a potential therapeutic strategy.

## Materials and methods

### Animal models

All procedures involving zebrafish followed the Animal (Scientific Procedures) Act 1986 under the Home Office Project Licence (PPL P254848FD). Adult animals were raised and maintained in the Biological Services Aquarium under careful monitoring at 28°C under a 14/10 hour light/dark regimen. Zebrafish embryos were maintained in Petri dishes of approximately 60 embryos in E3 media in light cycling incubators at 28°C until 5 days post-fertilisation (dpf). For adult assays (beyond 3 months of age), animals were sex and size matched for each assay. Prior to this age, sexing could not be performed but animals were size matched where possible.

*rnaset2^sh532^* mutant fish were previously generated using CRISPR/Cas9 genome editing technology in *Tg(mpeg1:mCherryCAAX)sh378* embryos with founders identified and bred to produce a stable line [4, 21]. Wild type and homozygous adults were identified using fin clip genotyping and isolated for breeding. For isolation of GFP-positive macrophages, *Tg(fms:GFP)sh377* animals were used [10].

### Zebrafish-to-zebrafish immune cell transplantation

For zebrafish-to-zebrafish transplantation, macrophages were isolated from whole kidney marrow using flow cytometry, before injection into microglia-depleted embryos via the systemic circulation.

To prepare grafts for transplantation, whole kidney marrow was isolated from ten transgenic adult fish with GFP-labelled macrophages *Tg(fms:GFP)sh377* and GFP-negative controls (nacre). Animals were culled by schedule 1 methods: first, animals were supplied with terminal anaesthesia in a tricaine bath (1.33g/L in aquarium water) followed by destruction of the brain. Culled animals were then secured on an agarose dissection plate with the abdominal cavity exposed via a deep incision along the ventral midline. To retrieve the kidney, internal organs were removed (including any egg debris) and the exposed kidney was peeled from the dorsal surface using forceps. Each kidney was then placed in 200μl of cold live sorting buffer (L15 media, 20% foetal bovine serum and 5mM EDTA) and mechanically separated using repeated pipetting by P1000 and P200 pipettes, to encourage dissociation of cells from connective tissue. Samples from each fish were kept separate, and on ice, until immediately before flow cytometry to prevent cross-reactivity or immune activation. Finally, samples were filtered through FACS tubes and pooled for sorting.

To isolate GFP-positive cells, samples were loaded into the BD FACSMelody^TM^ cell sorter (BD Biosciences) at 4°C. GFP-negative samples were sorted first to determine thresholds for GFP-fluorescence, and TO-PRO^TM^3 (ThermoFisher, R37170) was used to facilitate removal of dead cells. GFP-positive cells were sorted into an Eppendorf tube containing 500μl live sorting buffer and placed on ice before transplantation. Cell yields of approximately >500,000 cells were optimum for transplantation.

After FACS, GFP-labelled cells can become photobleached, making visualisation and quantification of transplanted cells challenging in host embryos for several days post-transplantation. To allow immediate visualisation, sorted cells were labelled with the fluorescent dye carboxyfluorescein succinimidyl ester (CFSE) by incubation in 1:10,000 CFSE dilution for 15 minutes at 26°C.

After graft preparation, transplanted cells were centrifuged with excess supernatant removed until graft contents could be transferred to microcentrifuge tubes. Cells were suspended in 1% polyvinylpyrrolidone (PVP) in live sorting buffer, before final centrifugation and resuspending at a concentration of approximately 20–30 cells per nanolitre.

Transplants were performed on 2dpf embryos with their endogenous microglia and macrophages labelled with mCherry (*Tg(mpeg1:mCherry CAAX)sh378*). Where required, these embryos had undergone microglia depletion via CRISPR/Cas9-mediated knockout of *irf8* (described below). On the day of transplantation, hosts were dechorionated, anaesthetised with 4.2% tricaine and lined up on a moist 28°C agarose plate for injection. Sorted cells were injected into the systemic circulation via the duct of Cuvier. Hosts were then rescued from anaesthesia in fresh E3 and returned to the incubator for recovery. As a negative control, approximately 60 embryos received a sham transplant containing only 1% PVP in live cell sorting buffer, without any cells present.

Following transplantation, embryos were sedated with 4.2% and screened using a Zeiss Axioscope to confirm the presence of GFP-positive cells. For longitudinal quantification of transplanted cells, positive embryos were screened at 3dpf and transferred to individual wells of 48-well plates for follow-up. For all other assays, embryos were screened at 5dpf and those with 10 or more GFP-positive cells in their brain were taken forward for subsequent assays. For raising until 8dpf, embryos were transferred to fresh Petri dishes containing E3 media and returned to the light cycling incubator. Media was changed daily, along with the addition of fresh food and removal of any unhealthy larvae for culling by schedule 1. For raising to adulthood, embryos were housed at densities of 15–20 animals per tank and fed twice daily to ensure healthy development.

### Microglia depletion by knockdown of irf8

To deplete microglia from embryonic brains, CRISPR/Cas9 genome editing was utilised to target interferon regulatory factor-8 (*irf8*) – a transcription factor essential for the development of macrophages and microglia [30]. To achieve maximum depletion, two guides targeting *irf8* expression (see **Table S1**) were injected at a final concentration of 50μM each by creating an injection mix of 0.5μl of each guide at 100μM, 1μl of 50μM tracer and 1μl Cas9 nuclease (New England Biolabs, MO386M). As a control, scrambled crRNA was used at 50μM in the place of *irf8*-targe ting guides. For each embryo, 2μl of injection mix was injected into the yolk sac at the single cell stage. Fish were screened for survival using light microscopy each day following injection, with unhealthy or dead embryos removed. To assess microglia depletion, the number of *mpeg:mCherry* cells in the brain were quantified at 5dpf using ZEISS Axio Scope.

### Whole mount TUNEL and immunohistochemistry

In order to assess the functionality of transplanted cells in *rnaset2* mutant embryos at 5dpf and 8dpf, TUNEL staining (to visualise uncleared apoptotic debris) and immunohistochemistry (to visualise expression of microglia-specific markers) were performed in parallel on the same samples.

Following screening to ensure transplanted cell engraftment and effective microglia depletion (where necessary), 5dpf and 8dpf embryos were immersion fixed in 4% PFA overnight at 4°C following terminal seda tion in concentrated tricaine. Following fixing, samples were rinsed with PBST and dehydrated with increasing concentrations of methanol (25% MeOH:75% PBS; 50% MeOH:50% PBS; 75% MeOH:25% PBS; 100% MeOH) and stored at –20°C until TUNEL assay and/or immunohistochemistry.

To quantify apoptosis, we utilised the terminal deoxynucleotidyl transferase (TdT)-mediated dUTP nick end labelling (TUNEL) assay – a standard protocol in the assessment of controlled cell death – using the commercially available Apoptag Kit, as per manufacturer instructions.

Following completion of the TUNEL, immunohistochemistry was performed. Embryos were incubated in primary antibody solution for 48 hours at 4°C at following concentrations: anti-4C4 mouse antibodies at 1:100 and anti-GFP chicken antibodies (GeneTex, GTX13970) at 1:500. Anti-4C4 primary was isolated from 7.4.C4 mouse hybridoma cells (Sigma, 92092321-1VL) as per manufacturer instructions.

Samples were then washed and incubated in the relevant secondary antibodies for 24 hours at 4°C.: Alexa Fluor^TM^ 647 goat anti-mouse (Thermo-Fisher Scientific, A-21235) and Alexa Fluor^TM^ 488 goat anti-chicken (Thermo-Fisher Scientific, A-11039).

Immunostained embryos were mounted in 0.1% low melting temperature agarose and imaged using Nikon W1 Spinning Disk. Each embryo was imaged using 20x magnification, with a Z-stack of 50 slices (2µm per slice). Maximum intensity projections generated using Fiji are shown throughout. Imaging and quantification were performed blinded to minimise bias. To establish the region of interest, the optic tectum was identified using a brightfield reference image. TUNEL counts and 4C4-GFP co-localisation were then counted manually using Fiji.

### Tissue clearing

Tissue clearing and immunohistochemistry methodology was adapted from Susaki *et al. Nature Protocols* 2015 [50] and Ferrero *et al. Cell Reports* 2018 [15].

For long-term assessment of transplant engraftment, successfully transplanted animals were raised according to standard zebrafish husbandry. Fish were checked weekly to observe any potential side effects of transplantation, and to maintain fish health. At the desired timepoint, fish were culled by decapitation with each head placed into 2ml 4% PFA and gently inverted to ensure the entire tissue was fully submerged, before leaving at 4°C overnight to ensure fixation of the tissue. Samples were then rinsed in autoclaved PBS and brains dissected in a Petri dish containing PBS placed on ice under a light microscope. Dissected brains were then gradually dehydrated using sequential 30-minute washes in methanol: 25% MeOH: 75% PBS; 50% MeOH: 50% PBS; 75% MeOH: 25% PBS; 100% MeOH). For long-term storage, samples were placed in fresh methanol and stored at –20°C to ensure preservation of tissue and quenching of fluorescent signal to minimise interference with subsequent staining.

CUBIC reagents were prepared according to Susaki *et al. Nature Protocols* 2015: CUBIC-1: 25% (by weight) urea (Sigma, 15604-1KG), 25% Quadrol (Sigma, 122262-1L), 15% Triton X-100 (Sigma, X100-500ML) in dH2O; CUBIC-2: 25% urea, 50% sucrose (Fisher, 10386100), 10% triethanolamine (Sigma, 90278-100ML) in dH2O [50].

At the time of clearing, samples were rehydrated from 100% methanol using serial MeOH dilutions in dH_2_O, as PBS is known to interfere with clearing via CUBIC-1. At this stage, samples <14dpf were taken forward for immunohistochemistry without any additional clearing, due to their small size and transparency. For samples 28dpf and older, brains were then submerged in 1ml 50:50 CUBIC-1:dH_2_O for up to six hours. Following full equilibration of the sample (confirmed by checking that the sample has sunk to bottom of the tube), 50:50 CUBIC-1:dH_2_O was removed and 1ml CUBIC-1 was added. Samples were then left to incubate at room temperature for 3 days.

After CUBIC-1 clearance, samples were washed three times with PBS, before being immersed in 1ml 50:50 CUBIC-2:PBS and incubated for up to 24 hours. 50:50 CUBIC-2:PBS was then replaced with CUBIC-2, and samples left to incubate for 3 days for 28dpf animals, and 6 days for 42dpf and older.

After clearance, samples were ready for immunohistochemistry as described above. Samples were then digested in 1:500 proteinase K for 2 hours at room temperature, before three five-minute washes with PBS. Primary antibody incubation was 48 hours at 4°C. Secondary antibody incubation was 24 hours at 4°C. Finally, samples were incubated in 1:5000 DAPI solution (in PBS) for 60–90 minutes at room temperature, followed by thorough washes. Once staining was complete, the samples were re-cleared with CUBIC (overnight incubation with 50:50 CUBIC-2:PBS and 3–6 days in CUBIC-2) to ensure full transparency for imaging.

Samples were mounted in CUBIC-2 and aligned so that the dorsal surface of the brain was flush against the coverslip, before imaging with the Nikon W1 spinning disk confocal microscope. For each sample, a 500μm Z-stack was taken with 10x or 20x lens, using the Z-spacing recommended by NIS Elements (0.9µm for 20x images, 2.5µm for 10x images). Large images were taken using the tiling function (3×2 grid) within NIS elements. Samples from each timepoint were imaged on the same day to ensure fair comparison between groups. Analysis was performed with Arivis as described below.

### Semi-automated quantification of cleared adult tissue

An automated pipeline was established using Arivis Vision 4D, which allows sophisticated segmentation of complex datasets. This quantification pipeline began by converting the ND2 file from the Spinning Disk into an Arivis compatible SIS file. The optic tectum was manually identified as the region of interest from each whole brain image, as processing the whole image was beyond the capabilities of the analysis computers available in the Wolfson Light Microscope facility. The optic tectum was chosen as a region of interest to allow alignment with the embryonic data sets from other transplant assays (TUNEL etc.), and because this area remained robustly intact after dissection, immunohistochemistry and clearing while other more peripheral regions (such as the olfactory bulb or hindbrain and spinal cord) were frequently damaged. To begin quantification, a closing morphology filter was applied in other to smooth the appearance of otherwise highly ramified and irregular microglia to create a more uniform structure that can be better identified by subsequent segmentation tools. The ‘blob finder’ segmentation tool was then performed – an analysis tool suited to finding irregular, rounded objects. The parameters of this blob finder segmentation were adjusted to most accurately capture the relevant cell population, using an average diameter of 30µm for objects in the GFP channel, and 15µm for 4C4-positive cells. Diameters were established by using manual measurement of cells in the raw image as a starting point, then adjusting the value until the segmentation most accurately mapped visible cells. The differing diameters set for each channel – despite labelling the same cell type – are attributed to the different cellular localisation of the marker proteins: *fms:GFP* being a cytoplasmic marker distributed throughout the cell, while the 4C4 antigen may be more discrete. Using these parameters, the blob finder segmentation was able to identify cells in three dimensions.

To remove any analysis artefacts, the resulting segments were filtered to exclude any objects with a volume larger than 5000µm^3^ or less than 300µm^3^. These values differ to those used for the Imaris segmentation – however, this is because Imaris requires spherical estimations of volume to provide a threshold, while Arivis is better able to calculate the volume of irregular objects and filter accordingly. At the final stage, objects were manually filtered by intensity to account for differing staining efficiencies between timepoints to ensure the final object count best reflected the true number of cells.

To establish the co-localisation of 4C4 and GFP signal, a parent-child analysis was performed to quantify the number of GFP-positive objects that contained a 4C4-positive object, with a minimum overlap of 20%. Data was then exported into an Excel spreadsheet and visualised with GraphPad Prism.

### Larval RNA extraction and cDNA synthesis

Prior to RNA extraction, embryos (5–8dpf) were culled by terminal anaesthesia in concentrated tricaine, before being transferred to Eppendorf tubes and the addition of 500µl Trizol reagent (Invitrogen, 15596026). Once in the Trizol reagent, culled embryos were homogenised using a handheld homogenizer (T 10 basic ULTRA-TURRAX, IKA Dispersers, product no. 0003737002). Following complete homogenisation, 100µl chloroform was added and the sample was mixed vigorously. After 2–3 minutes at room temperature, samples were centrifuged at 12,000g for 30 minutes at 4°C. The upper aqueous phase was removed (200µl) and transferred to a clean Eppendorf with 200µl isopropanol, before leaving overnight at –20°C. The following morning, samples were again centrifuged at 12,000g for 30 minutes at 4°C before the supernatant was removed and pellet washed with 1000µl ethanol. Samples were centrifuged at 12,000g for 15 minutes at 4°C (repeated as needed) until all supernatant could be removed and the pellet left to air-dry for 10 minutes. Finally, the resultant pellet was dissolved in 20µl nuclease-free water to be used for cDNA synthesis.

cDNA was synthesised utilising Superscript II according to manufacturer’s instructions. Briefly, 2µg RNA was transferred to a microcentrifuge tube with 1µl oligo(dT), 1µl dNTP mix and nuclease-free water to 10µl before incubating at 65°C for 5 minutes. Samples were briefly chilled on ice, before the addition of 4µl 5x First Strand Buffer, 2µl DTT, 1µl RNase-OUT RNase inhibitor and 1µl Superscript II reverse transcriptase. The mixture was incubated to 42°C for 50 minutes before inactivation at 70°C for 15 minutes. The resultant cDNA was diluted 1:20 for qPCR.

### qRT-PCR

qRT-PCR was performed to assess relative gene expression. Reaction mixtures were as follows: 5μl SYBR green, 2μl milli Q water, 2μl cDNA, 0.5μl forward primer and 0.5μl reverse primer (both 10µm). The qRT-PCR reaction was run on a CFX96 Bio-Rad machine as follows: Step 1: 95°C for 2 minutes, Step 2: 95°C for 10 seconds, Step 3: 57°C for 30 seconds, Step 4: 72°C for 25 seconds, Step 5: 95°C for 30 seconds, Step 6: 65°C for 10 seconds, Step 7: 95°C for 20 seconds with Steps 2–4 repeated 39 times and increment of 0.2°C every 10 seconds between Steps 6 and 7. As previously published, ef1α was used as a reference gene for each sample [21]. Each reaction was performed in triplicate, with the mean Cq value for each reaction used to determine relative expression as below:

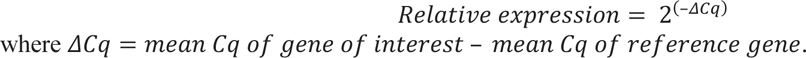

### Larval free-swimming analysis

In order to assess larval free-swimming behaviour, embryos were raised to 8dpf under approved individual study plan. Briefly, animals were raised in Petri dishes – 12 embryos per dish – containing E3 media which was refreshed daily by the personal licence holder. Embryos were fed daily, using a Pasteur pipette to ensure uniform distribution of food between plates. On the morning of the experiment, embryos were transferred to 24 well plates (one embryo per well) without anaesthetising, with 500µl clear E3 per well. Plate layout was alternated between experimental groups to ensure no location-specific effects – and allowed to habituate for at least 4 hours. Swimming distance was recorded across a 20–60-minute period, with alternating light-dark cycles of 10 minutes. Outliers were manually excluded based upon movement analysis trace automatically generated by the Zebralab software.

### Juvenile free-swimming analysis

For adult swimming behaviour analysis, fish were raised as normal to juvenile/adult stages. Animals were transferred to the behavioural facility on the morning of recording and allowed an initial 30-minute habituation period to recover from any stress when transferring between rooms. For recording, each adult was placed in an individual 0.7L tank (Tecniplast) containing 200ml aquarium water. The walls of each tank were covered with white paper to ensure the animals were unable to interact or be distracted by their surroundings.

After transfer to the behavioural tanks, fish were given a further 10 minutes to habituate to their environment, before recording for a period of 10 minutes. Recording was performed using the Basler GenICam camera and tracking performed using EthoVision XT15 (advanced detection settings with dynamic subtraction). Total distance swum was extracted from EthoVision software.

Following recording, animals were either culled by schedule 1 or by decapitation (regulated K) if their brains were required for further experiments. Animals were size matched throughout. At the 4wpf timepoint, animal sex could not be determined due to their small size and early development.

### Survival analysis

Survival at 4wpf was defined as the absence of any observable sickness behaviours. Animals were routinely monitored, and any animal with inability to swim to feed was humanely culled.

### RNA sequencing

For RNA sequencing, 4wpf transplanted *rnaset2* mutants and sham controls were culled and brains dissected as previously described. Dissected brains were placed immediately into liquid nitrogen to snap freeze the tissue and preserve RNA integrity for extraction at a later timepoint. RNA was then extracted using Trizol as described above, except for the final step in which the resulting RNA pellet was dissolved in 100µl nuclease-free water (rather than 10µl) to aid column purification.

The RNeasy MinElute Cleanup Kit (Qiagen, cat. no. 74204) was used to purify RNA for sequencing. According to manufacturer’s instructions, the dissolved RNA was mixed with 350µl buffer RLT (from kit) before the addition of 250µl 100% ethanol to promote binding of the RNA to a silica membrane. This mixture was then transferred to an RNeasy MinElute spin column (from kit) and centrifuged for 30 seconds at 10,000 rpm. The membrane was then washed with 350µl buffer RW1 to remove contaminants, and span again 30 seconds at 10,000 rpm. On recommendation by our sequencing provider, we then added 10µl DNaseI mixed with 70µl RDD buffer to each column and incubated at room temperature for 15 minutes to remove any gDNA contamination. Following DNaseI treatment, samples were washed again in buffer RW1, then again with 500µl of buffer RPE and finally with 500µl 80% ethanol. After drying, 14µl nuclease-free water was then added to the centre of the silica membrane to elute the RNA, which was then stored at –80°C until quality control.

Quality control was performed by The Genomics Laboratory at the University of York, using the Agilent BioAnalyzer 2100. Samples were first assessed for quality using the RNA Integrity Number (RIN) given by the BioAnalyzer, which measures the ratio of each ribosomal subunit along with potential degradation products to give a measure of RNA quality. Samples with a RIN exceeding 7.0 were taken forward for sequencing.

Library preparation was performed by The Genomics Laboratory at the University of York, using the NEBNext Ultra II Directional Library prep kit for Illumina in conjunction with the NEBNext® Poly(A) mRNA Magnetic Isolation Module and unique dual indices (New England Biolabs), according to the manufacturer’s instructions. Libraries were pooled at equimolar ratios and sent for paired end 150 base sequencing at Azenta Life Sciences on an Illumina NovaSeq 6000 instrument.

Reads (data accession number PRJNA1047321) were trimmed using Cutadapt v3.4 [33], then mapped to the GRCz11 genome using Spliced Transcripts Alignment to a Reference (STAR) v2.7.10b [11]. Counts were generated for each gene using htseq-count v2.0 [43]. Differential expression analysis was performed using DESeq2 [31] using three-way comparisons between wild type sham, *rnaset2* sham and rnaset2 transplanted samples (with separate analyses for microglia-depleted and non-depleted groups). Pathway analysis was performed using g:Profiler (https://biit.cs.ut.ee/gprofiler/gost) [28]. For GSEA, genes were ranked according to their Wald statistic results from the differential expression analysis. The ranked list of genes was then used for a Gene Set Enrichment Analysis using the package *fgsea* v1.28.0 and the results were plotted using the package *ggplot2* v3.4.4. in R v4.3.2.

### Statistical analysis

All statistical analysis was performed using GraphPad Prism throughout. For pairwise comparisons, data was entered using a column (two samples, one variable only) and analysed using Mann Whitney test. For comparisons between more than two samples, data was entered using a grouped table and analysed using Kruskal-Wallis test with Dunn’s multiple comparisons. For survival analysis, log rank Mantel-Cox test with Bonferroni’s multiple comparison correction was used. Exact *p* values are stated throughout.

For larval experiments, assays were repeated three times using different batches of larvae born on different dates, with the number of biological replicates and n (experimental unit) number stated for each experiment in figure legends unless otherwise stated. For juvenile experiments, each animal represents an experimental unit (n stated in figure legend).

For RNA sequencing, significance testing was performed using Wald tests with Benjamini-Hochberg p value correction, as previously described [31].

## Results

### Macrophage transplantation successfully replaces microglia in wild type zebrafish

We have previously demonstrated that *rnaset2* microglia are deficient in their ability to clear apoptotic debris - leading to a downstream neuroinflammatory response [21]. As such, we sought to replace these dysfunctional microglia using transplantation of healthy macrophages. Although HSC transplantation is used clinically in hereditary leukodystrophies and may act to replace microglia, transplanted macrophages may represent a quicker route to repopulation of the brain – minimising the need for HSC engraftment and instead migrating directly to the central nervous system (CNS). To this end, healthy macrophages were isolated from whole kidney marrow (WKM) from *Tg(fms:GFP)* adult fish and purified by FACS. Purified macrophages were injected at 2 days post-fertilisation (dpf) into embryos with endogenous macrophages depleted by CRISPR/Cas9 knockout of the transcription factor *irf8* (**Figure 1a**). Transplanted macrophages were able to robustly engraft within host embryos and reach the brain within three days of transplant in microglia-depleted animals (**Figure 1b, c)**. The numbers of transplanted cells in the brain increased in the days post-transplant, with timelapse imaging confirming that transplanted macrophages were able to divide following engraftment in the brain (**Figure 1c, supplementary video 1**). Conversely, transplantation of CD41-positive haematopoietic stem cells was less efficient therefore confirming the suitability of WKM-derived macrophages as a graft resulting in a more direct repopulation of the microglial niche (**Figure S1**). Depleting the host of endogenous microglia using *irf8* crRNA CRISPR/Cas9 injection promoted transplanted cell engraftment, highlighting the importance of emptying the niche before transplantation (**Figure S1a,b).**

**Figure 1.**
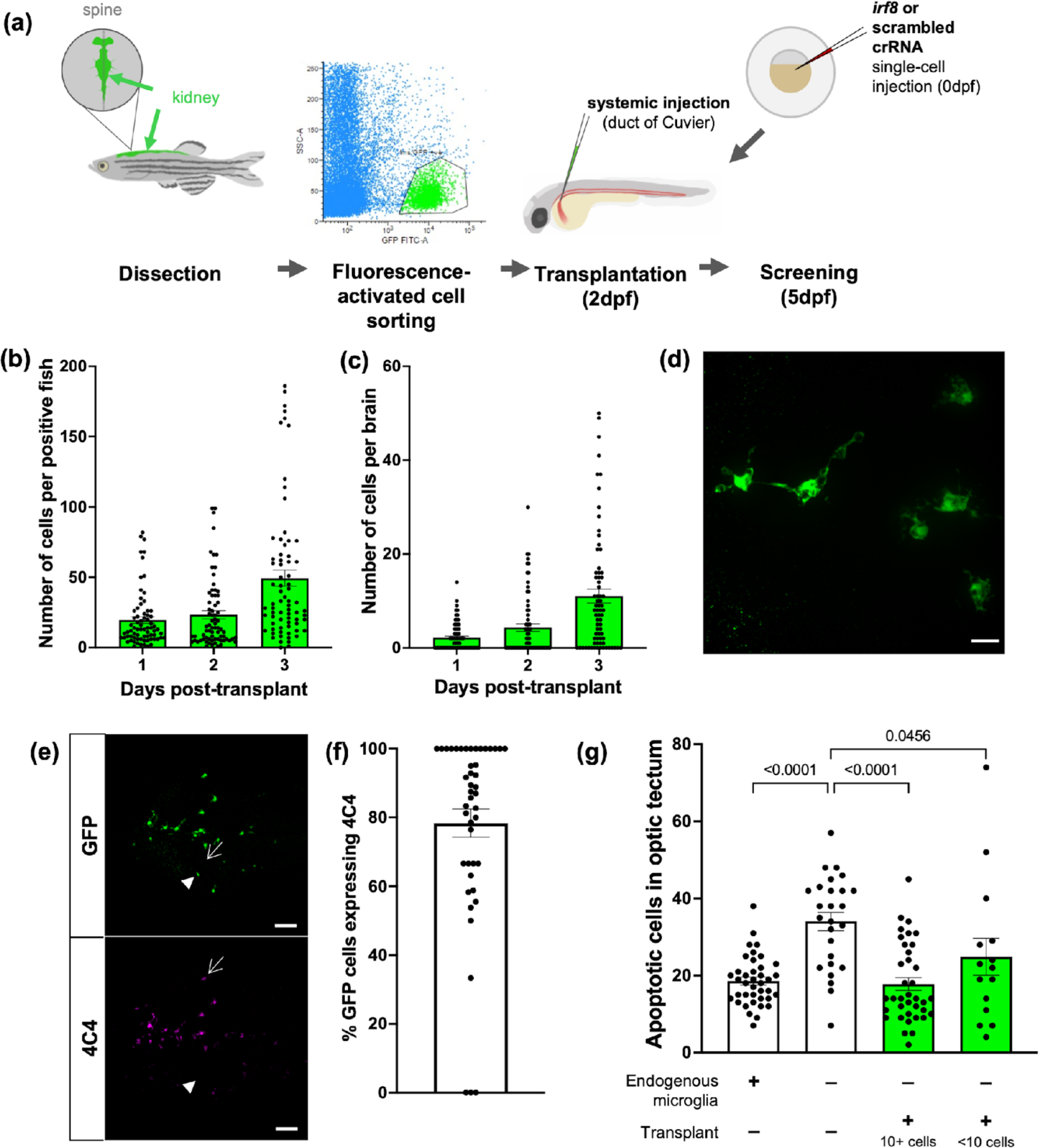
Macrophage transplantation successfully replaces microglia in wild type zebrafish. **a.** Transplants were performed by dissecting kidneys from *Tg(fms:GFP)* adult donors and isolating GFP-positive cells using fluorescence-activated cell sorting. Cells were injected into 2dpf hosts via duct of Cuvier. Hosts had previously undergone microglia depletion via knockout of *irf8* or were non-depleted with a scrambled control. **b, c.** The number of *fms:GFP* cells per successfully transplanted animal increased throughout the whole body **(b)** and within the brain **(c)** within 3 days post-transplant. **d.** Confocal imaging reveals that transplant-derived cells show a branched, microglia-like morphology. Scale bar represents 17µm. **e, f.** Transplant-derived cells express the microglia-specific marker 4C4 at 3 days post-transplant (5dpf). Pointed arrow indicates cell positive for both GFP and 4C4, closed arrow indicates GFP-positive cell that does not co-localize with 4C4. Three biological replicates, n=45. Scale bar represents 75µm. **g.** TUNEL staining reveals that transplantation can rescue the number of uncleared apoptotic cells in microglia-depleted brains in a dose-dependent manner. Kruskal-Wallis test with Dunn’s multiple comparisons. Four biological replicates, n=15-39.

Transplanted macrophages displayed a highly microglia-like morphology, with multiple branches surveilling the CNS environment (**Figure 1d**). As such, we sought to investigate whether these cells could adopt aspects of the microglial phenotype following engraftment. Using colocalisation with the microglia marker 4C4 [45], we showed that up to 80% of transplant-derived cells express microglia-specific proteins - suggesting that these WKM macrophages are able to adapt to their host environment (**Figure 1e,f**). Crucially, transplantation was associated with reduced abundance of apoptotic cells in a dose-dependent manner, demonstrating the ability of transplant-derived cells to phagocytose dying cells within the CNS (**Figure 1g**). Together, these data suggest that transplanted cells are able to reprogramme as microglia and undertake tissue-resident functions.

### Macrophage transplantation reduces early neuropathology in rnaset2 mutants

After demonstrating that transplanted macrophages can repopulate host brains in microglia-depleted wild type animals, we next sought to investigate whether transplantation had comparable efficacy in *rnaset2* mutants. As *rnaset2* microglia are deficient in their ability to clear apoptotic cells accumulating during development – known to be one of the key drivers of microglial infiltration into the developing brain – we investigated whether endogenous microglia depletion was necessary for complete engraftment [6]. Interestingly, we found significantly more transplant-derived cells in the brains of microglia-depleted hosts compared to non-depleted controls, suggesting that transplanted macrophages compete with dysfunctional endogenous microglia cells for brain engraftment, as found in WT brains (**Sup Figure 1 and Sup Figure 2**). However, we also noticed that a greater proportion of transplanted cells expressed the microglia-specific marker 4C4 in microglia-depleted animals – suggesting that these cells are able to take on a microglial-like phenotype in *rnaset2* mutants when endogenous microglia are absent, but retain aspects of their macrophage identity when host microglia are present (**Figure 2a,b**). This failure to express microglia-specific markers was found in transplanted cells in non-depleted brains both 3- and 5-days post-transplant – suggesting this is not due to a delay in cells entering the brain where they may undergo reprogramming, but rather due to an overall failure to adopt a microglia-like phenotype.

**Figure 2.**
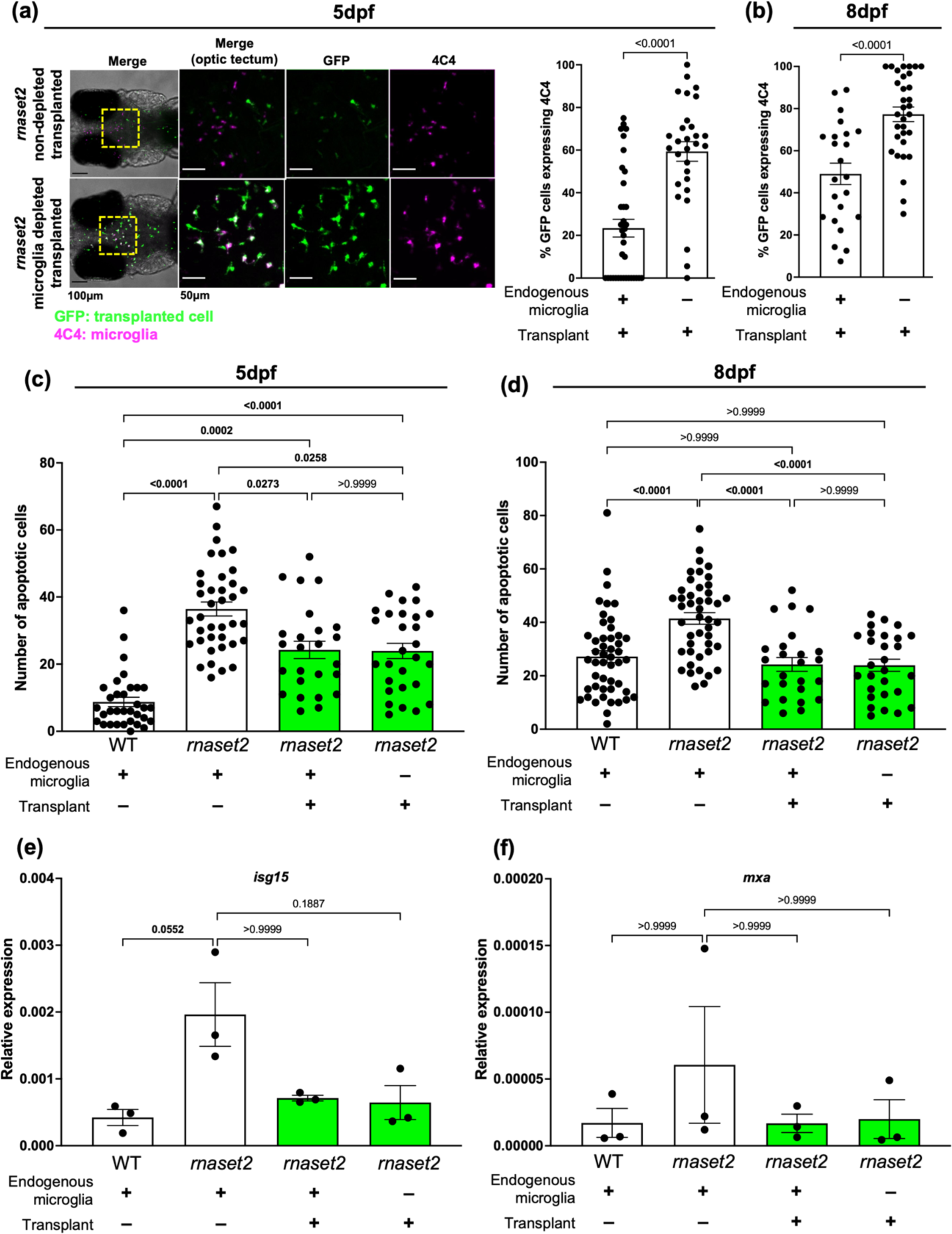
Macrophage transplantation reduces early neuropathology in rnaset2 mutants. **a,b**. Quantification of 4C4-expression of transplant-derived cells in microglia-depleted versus non-depleted *rnaset2* hosts at 5– **(a)** and 8dpf **(b)** reveals reduced expression of 4C4 by transplanted cells in non-depleted hosts. Mann Whitney test. 3 biological replicates, n=29–37 **(a).** 3 biological replicates, n=23–32 **(b). c,d.** Macrophage transplantation reduces the number of uncleared apoptotic cells in the optic tectum of 5– **(c)** and 8dpf **(d)** *rnaset2* mutants. Kruskal-Wallis test with Dunn’s multiple comparisons. 4 biological replicates, n=25–38 **(a)**. 3 biological replicates, n=25–48 **(b)**. **c.** Representative images of TUNEL counts in the optic tectum (red dashed line). Scale bar represents 100µm. **e,f.** qPCR reveals that *isg15* **(e)** *and mxa* **(f)** expression is reduced in transplanted animals, relative to wild type controls. Kruskal-Wallis test with Dunn’s multiple comparisons. Three biological replicates, 15 embryos pooled per replicate.

After demonstrating that transplanted macrophages are able to engraft in *rnaset2* embryos, we next sought to investigate whether this intervention could rescue neuropathology observed in our disease model. As a failure to digest dying cells during development is thought to be one of the initiating factors in *rnaset2* pathology, we first investigated the ability of transplanted macrophages to clear apoptotic debris in *rnaset2* mutants. Indeed, TUNEL staining revealed that transplanted animals exhibited significantly fewer uncleared apoptotic cells at 5– and 8dpf, with a complete rescue to wild type levels at 8dpf (**Figure 2c,d**). Interestingly, this clearance of apoptotic cells was comparable across transplanted animals regardless of endogenous microglia depletion at both ages. We had previously hypothesised that this bottleneck of apoptotic cell digestion may trigger downstream neuropathology - in particular, the antiviral response which is prominently upregulated in mutants during larval and adult stages. As such, we next investigated the impact of transplantation on antiviral responses in the heads of 8dpf animals. qPCR revealed an approximately four-fold downregulation of the interferon response gene *isg15* in transplanted mutants – a correlate of the antiviral response significantly upregulated in the *rnaset2* mutants (**Figure 2e**). A similar pattern of rescue was observed for other antiviral genes, including *mxa* (**Figure 2f**). This normalisation of the antiviral response could not be explained by elevated levels of functional, transplant-derived rnaset2, as *rnaset2* transcript levels remained downregulated in transplanted mutants compared to wild type controls (**Supplementary Figure 3**). Therefore, these findings suggest that the presence of phagocytosis-competent macrophages in the CNS can stabilise the antiviral cascade in *rnaset2* brains.

### Long-lasting transplanted macrophages rescue neuroinflammation beyond embryonic stages

To assess whether transplanted cells persisted in the brain to provide long lasting therapeutic effects, we used tissue clearing followed by immunohistochemistry on brains from WT transplanted animals to quantify endogenous versus transplanted cell number over time. We found that transplant-derived cells initially expand – peaking at 4 weeks post-fertilisation (wpf) – but were no longer detectable by 14wpf (**Figure 3a–c**). This reduction in transplanted cell number appeared to be accompanied by a gradual reduction in the percentage of remaining transplant-derived cells which express the microglial-specific marker 4C4 (**Figure 3d**). Interestingly, like in microglia-depleted wild type hosts, we saw a robust engraftment of transplanted cells maintained in microglia-depleted *rnaset2* brains until 4wpf, which was also cleared by 14wpf (**Figure 3e-g**). However, in non-depleted *rnaset2* mutants, transplanted cells were largely absent by 4wpf - suggesting that these cells are cleared more quickly when host microglia are present. Together, these data suggest that transplant-derived cells are able to maintain engraftment for the first few months of life in hosts lacking endogenous microglia during early development, regardless of genotype. Therefore, in order to investigate the impact of transplantation on *rnaset2* pathology beyond embryonic stages, we utilised microglia-depleted 4wpf hosts for our subsequent experiments.

**Figure 3.**
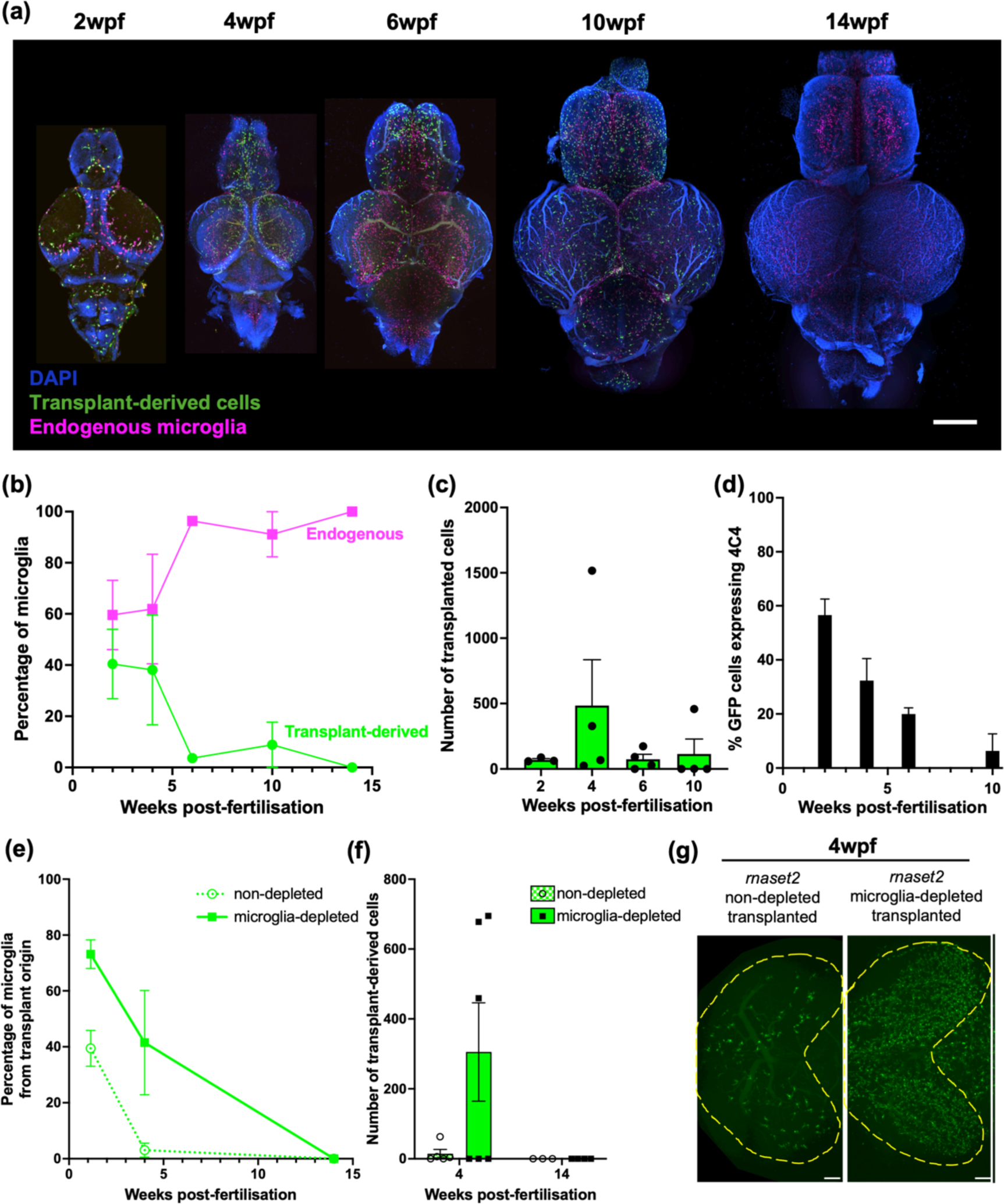
Transplanted cells persist in host brains throughout juvenile stages. **a.** Tissue clearing and immunohistochemistry reveals that transplant-derived cells persist in host brains until 10wpf but are cleared by 14wpf. Scale bar represents 400µm. **b.** Percentage of microglia from transplant origin gradually decreases in host brains until 14wpf. **c.** The number of transplant-derived cells peaks at 4wpf and rapidly decreases thereafter. **d.** Percentage of transplant-derived cells expressing microglia marker 4C4 steadily decreases from 2wpf. Three to four biological replicates per timepoint. **e, f.** Tissue clearing and immunohistochemistry reveal reduced numbers of transplant-derived cells in non-depleted *rnaset2* animals compared with microglia-depleted siblings. Cells are no longer visible in both host groups by 14wpf. **g**. Representative image of transplant-derived cell engraftment at 4wpf in non-depleted and microglia-depleted *rnaset2* hosts. Yellow line indicates optic tectum outline. Scale bar represents 100µm.

We performed RNA sequencing on bulk brains from *rnaset2* microglia-depleted transplanted animals compared with microglia-depleted sham controls (n=4 brains per group) **(Figure 4a–c)**. We identified significantly enriched pathways in *rnaset2* microglia-depleted transplanted animals compared with *rnaset2* microglia-depleted sham control (**Figure 4c**), with a significant rescue in antiviral immune response pathways by gene set enrichment analysis (GSEA) (**Figure 4d)**. Antiviral immune pathways, such as ‘Response to virus’ and ‘ISG15 antiviral mechanisms’, were restored to WT in transplanted animals (**Figure 4e,f, Sup Figure 4**). Interestingly, immune-related pathways were not enriched in *rnaset2* non-depleted transplanted animals compared with non-depleted sham controls, likely due to a lack of transplant-derived cells in host brains at this timepoint (n=3 brains per group) (**Sup Figure 5, 6**). Nonetheless, this suggests that adding phagocytic competent healthy macrophages at early stages in brain development can rescue the brain-wide immune response, specifically suppressing the antiviral immune response, in *rnaset2* mutants.

**Figure 4.**
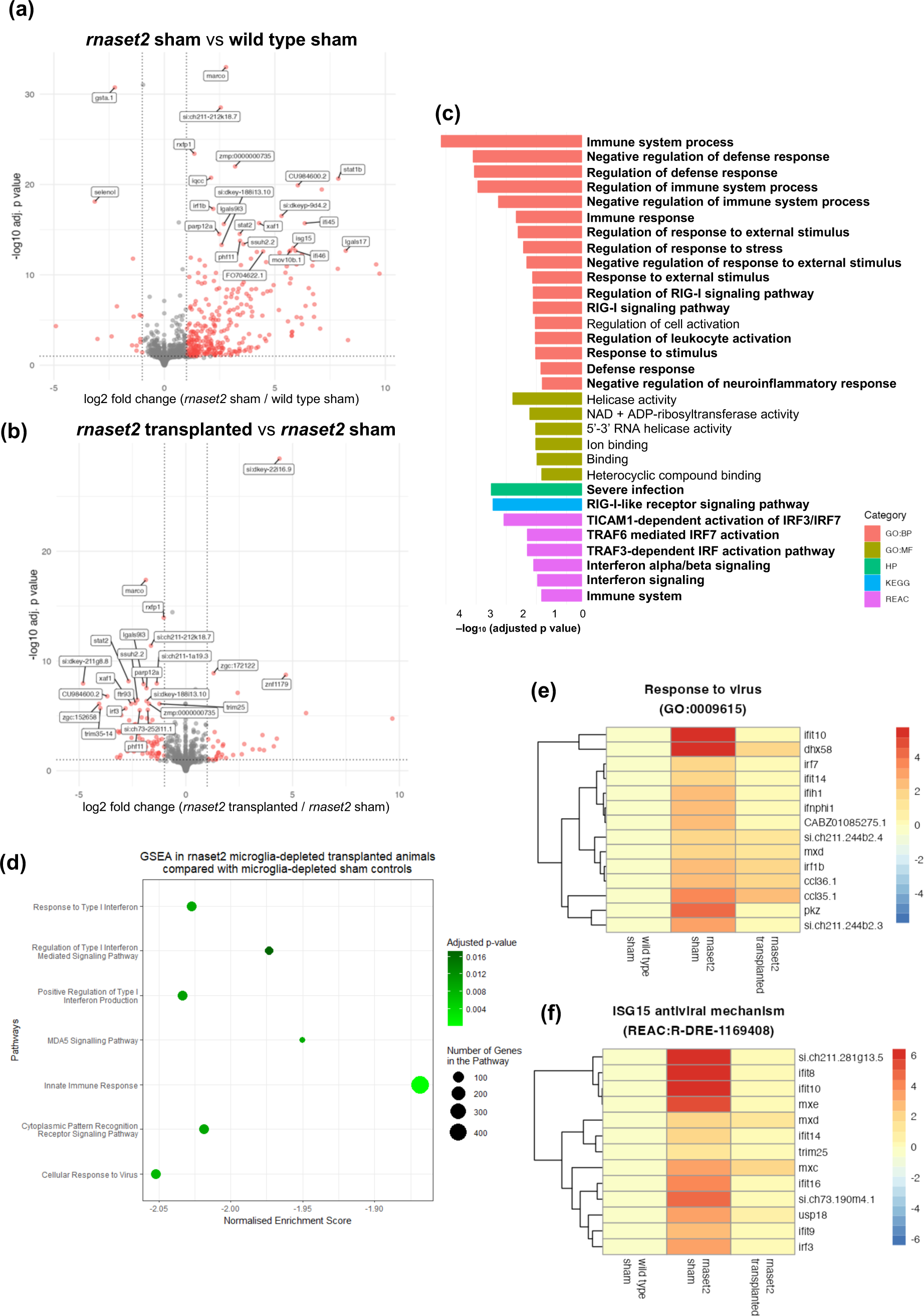
RNA sequencing reveals that microglia replacement recues antiviral immune response in 4wpf *rnaset2* mutants. **a, b.** Volcano plot of differentially expressed genes between *rnaset2* sham versus wild type sham **(a)** and *rnaset2* transplanted versus *rnaset2* sham groups **(b)**. Significantly differentially regulated genes are shown in red. The top 25 differentially expressed genes are annotated. **c.** Gene ontology (GO) plot showing significantly downregulated pathways in *rnaset2* microglia-depleted transplanted animals compared with microglia-depleted sham controls, identified by gProfiler. Pathways relating to immune and antiviral response are indicated in bold. **d.** GSEA in *rnaset2* microglia-depleted transplanted animals compared with *rnaset2* microglia-depleted sham controls. **e, f**. Heatmap of genes belonging to response to virus **(e)** and ISG15 antiviral mechanism **(f)** GO pathways for all genes significantly upregulated in *rnaset2* sham animals. Fold change relative to wild type sham is indicated by colour, with red indicating higher expression. All data shown corresponds to microglia-depleted animals. See also Figure S4.

### Transplantation rescues rnaset2 motor impairment and survival beyond embryonic stages

After demonstrating that transplantation rescued many of the hallmarks of *rnaset2* neuropathology, we next sought to investigate whether these changes in pathology translated to altered disease behaviour. Larval swimming analysis revealed that transplanted mutants swam greater distances over a 20 minute period at 8dpf compared to non-transplanted controls, which themselves were hypoactive relative to wild type (**Figure 5a,b**). This rescue was mirrored at 4wpf, with transplanted, microglia-depleted *rnaset2* mutants showing greater motor activity over a 10-minute period of free-swimming, compared with sham controls (Figure 5c). Notably, at this timepoint, non-depleted transplanted *rnaset2* animals were not assayed due to a failure of persistent engraftment in these animals by 4wpf (see **Figure 3e,f**). Interestingly, in microglia-depleted transplanted hosts, the extent of motor recovery was similar between juveniles which had persistent cell engraftment and those which did not – suggesting that there may be some residual benefits of transplantation shortly after these cells disappear from the brain. Strikingly, this behavioural improvement appeared to be accompanied by an increase in survival of transplanted *rnaset2* mutants at 4wpf relative to microglia-depleted sham controls. As such, macrophage transplantation has therapeutic benefits beyond embryonic stages lasting into juvenile stages in *rnaset2* mutants.

**Figure 5.**
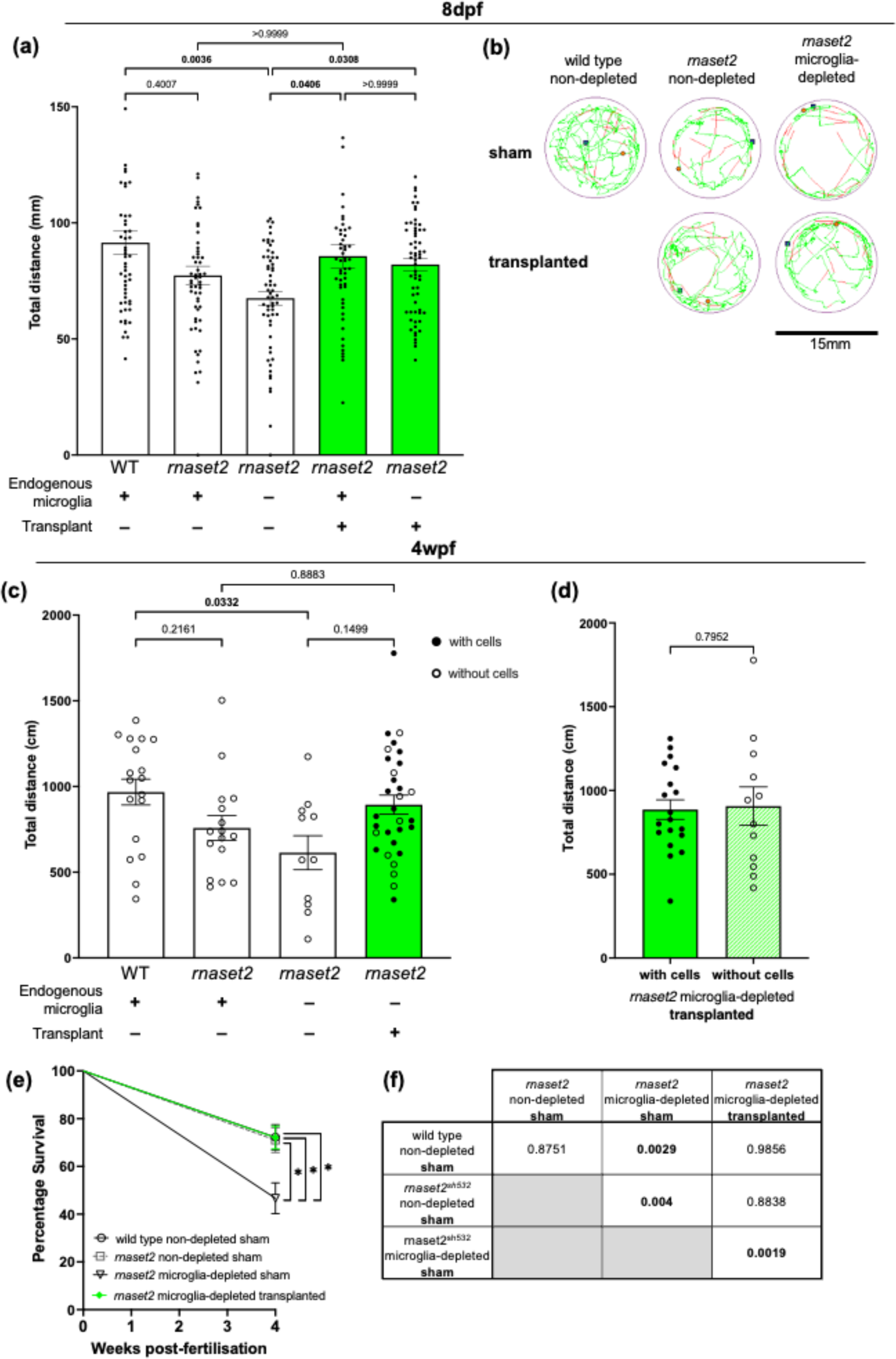
Transplantation rescues *rnaset2* mutant swimming behaviour and survival. **a.** Live tracking of larval swimming behaviour reveals that transplanted *rnaset2* mutants swim greater distances than non-transplanted controls. Kruskal-Wallis test with Dunn’s multiple comparisons. Three biological replicates, n=52–60. **b.** Representative traces of free swimming behaviour of zebrafish larvae. Green line represents slow movements (3.0–6.6mm/s), red line represents fast movements (>6.6mm/s). **c**. Tracking of 4wpf juvenile behaviour reveals that transplantation restores *rnaset2* swimming behaviour to wild type levels. Kruskal-Wallis test with Dunn’s multiple comparisons, n=11–31. **d.** Swimming behaviour was comparable between *rnaset2* microglia-depleted transplanted animals that had persistent engraftment of transplanted cells and those which did not. Mann-Whitney U test, n=12–19. **e, f.** Percentage survival of transplanted *rnaset2* mutants is greater than microglia-depleted sham controls at 4wpf. Pairwise comparisons and corresponding p values shown in **(f)**, bold indicates significance. Log rank Mantel-Cox test with Bonferroni’s multiple comparison correction (Bonferroni-corrected threshold: p<0.00833; family-wise significance threshold: P<0.05). Three biological replicates, n=60–86.

## Discussion

In this study, we have demonstrated that replacement of diseased microglia with healthy macrophages can rescue *rnaset2* pathology at a molecular, cellular and behavioural level. We have demonstrated that transplanted macrophages can reach embryonic brains, adopt a microglia-like phenotype and maintain engraftment for four to ten weeks post-transplantation. Transplanted macrophages compete with host microglia for filling of the microglial niche, even when host microglia are deficient in their ability to clear inflammatory debris in the brain. Nonetheless, the presence of transplanted cells was able to normalise the antiviral response, clear apoptotic debris and rescue the behavioural phenotypes shown by *rnaset2* mutants. RNA sequencing confirmed that macrophage transplantation rescues antiviral responses in *rnaset2* mutants at 4wpf juvenile stages. As such, this work supports the hypothesis that supplementation of healthy microglia improves aspects of pathology in RNASET2-deficient leukodystrophy, and that therapeutic targeting of the microglia may represent a future avenue for the development of novel treatments.

We have previously demonstrated that microglia-specific rescue of *rnaset2* expression can restore aspects of microglial function in *rnaset2* mutants – improving clearance of developmental apoptosis and restoring normal microglial morphology [21]. The current study extends this finding – showing that the presence of healthy phagocytes in the brain can not only clear apoptotic debris but rescue the neuroinflammatory response and restore motor function through larval and juvenile stages. Indeed, multiple other studies have supported the finding that microglial replacement can improve aspects of pathology – including behavioural rescue – across various neurodegenerative diseases [44, 47]. Additionally, treatment of neurological disorders with a cellular – rather than gene-based – therapy is particularly appealing in the context of microglial dysfunction, as many viral vehicles for gene delivery fail to efficiently target microglia at a whole-brain level [19, 22]. Our study therefore identifies a cellular mechanism whereby deficient microglia drive pathology in *rnaset2* mutants and supports a cellular approach to targeting microglia dysfunction in RNASET2-deficient leukodystrophy, beyond the genetic strategies previously explored.

Our data also evidence the ability of whole kidney marrow-derived macrophages to infiltrate the brain, divide and express microglia-specific markers. In the zebrafish, these cells are most similar to human bone marrow-derived macrophages and so may mimic peripheral blood cells following transplant. These cells have previously been demonstrated to infiltrate the brain following microglia depletion, during periods of inflammation or during embryogenesis (before the formation of the blood brain barrier) [9]. Due to the early timepoint at which our transplants were performed in the *rnaset2* mutant embryos, it is likely that all three of these factors contributed to successful engraftment in our intervention. However, the true extent to which transplant-derived macrophages become microglia-like remains unclear. Previous studies have demonstrated that, although macrophages derived from bone marrow and peripheral blood were able to engraft within the brain and express some microglia-specific markers, these transplant-derived cells remained transcriptionally distinct to host microglia even following robust engraftment [1, 52]. As such, it seems our results mirror the findings that transplant-derived macrophages can adapt to their new CNS environment and adopt key microglial phenotypes – but that they may remain distinct from their endogenous counterparts.

It is possible that the potential failure of these HSC-derived macrophages to fully adopt a microglial identity contributes to their lack of longevity in the host brain. However, it is interesting to note that there are two distinct populations of microglia in the zebrafish throughout development – with embryonic microglia derived from yolk sac progenitors gradually replaced throughout juvenile stages by a distinct population of adult HSC-derived macrophages [15]. This replacement is thought to be dependent on the arrival of the adult kidney marrow-derived second wave – with embryonic microglia persisting when HSC-derived microglia are genetically depleted [16]. Interestingly, this clearance of embryonic microglia follows a similar time course to that seen with our transplant-derived cells – despite these whole kidney marrow-derived cells being more similar in ontogeny to the HSC-derived adult microglia than the yolk sac embryonic progenitors. As such, it is possible that our transplant-derived cells undergo such extensive reprogramming following engraftment that they are recognised as self and follow the same trajectory as host embryonic microglia. It remains unclear whether this gradual decrease in transplant-derived cell number is due to cell intrinsic processes (such as cell death) or interactions with infiltrating host adult microglia. Transplantation of our macrophage graft into animals lacking this adult HSC-derived microglia population could, therefore, result in life-long engraftment of transplant-derived cells in the brain and potentiate their therapeutic effects.

It is interesting to note that, in order to achieve robust engraftment in *rnaset2* mutants, transient depletion of the embryonic microglial niche via knockout of *irf8* – a transcription factor required for the development of early microglia and macrophages – was required. We had initially speculated that the failure of rnaset2-deficient microglia to clear developmental apoptosis (among the first triggers to attract circulating monocytes to the CNS) would be sufficient for recruitment of transplanted macrophages to the brain [6]. However, without depletion of the embryonic microglial niche, the number of transplant-derived cells engrafted within host brains and the expression of microglia markers by these cells was much poorer than those with endogenous microglia depletion. This may have clinical implications, suggesting that myeloablation might be required in the context of transplantation in humans. However, despite these differences, the extent of pathology rescue was comparable between depleted and non-depleted hosts during larval stages, in terms of clearance of apoptotic debris, normalisation of the antiviral response and behavioural recovery. As such, it seems the presence of any additional healthy phagocytes in the brain can have benefits in *rnaset2* larvae, regardless of microglial identity. Additionally, even in microglia-depleted hosts, the mean number of transplant-derived cells remained much lower than the number of microglia which might be expected in a healthy age-matched embryo – suggesting transplant-derived cells may be particularly potent in their ability to clear neuropathology. Previous work has demonstrated that circulation-derived myeloid cells have an enhanced phagocytic ability relative to endogenous microglia [30]. Therefore, it seems that transplanted macrophages may be well suited to clearing apoptotic debris and restoring neuroinflammatory response regardless of their expression of microglial markers in *rnaset2* hosts.

Microglia replacement was sufficient to rescue the pathological antiviral and behavioural phenotypes of *rnaset2* mutants - however, future work is needed to assess the impact of this intervention of myelin structure and function. By 8dpf (the earliest timepoint used in this study), zebrafish show robust myelination which has been demonstrated to impact some behavioural functions [32, 41]. As such, our finding that microglia replacement rescues *rnaset2* locomotion presents two possible explanations. The first is that transplantation supports myelin integrity in *rnaset2* mutants. Such a hypothesis could be explored in future studies using myelin- and oligodendrocyte transgenic report lines, electron microscopy and MRI to investigate white matter function on a broader scale [20, 41]. An alternative hypothesis is that *rnaset2* locomotion defects are independent of myelination state at the ages investigated. While myelination has been linked to time-sensitive escape responses following startle stimuli in the zebrafish, a multitude of other factors can impact free swimming behaviours, including neuroinflammation and viral infection [8, 35]. Therefore, the interplay between transplant-derived cells, oligodendrocytes and myelination remains unclear. Nonetheless, the rescue of locomotion in transplanted animals suggests that the microglia are an attractive target for the development of future interventions in RNaseT2-deficient leukodystrophy.

In summary, our study highlights a cellular mechanism whereby microglia are the drivers of RNASET2-deficient neuropathology and suggests microglia-directed approaches may have therapeutic benefits in leukodystrophy. We demonstrate that adult whole kidney marrow macrophages can engraft in host brains and undertake microglial phenotypes, both in wild type animals and our disease model. Restoring microglia function by macrophage transplantation was sufficient to rescue brain-wide autoimmune phenotypes and restore locomotor activity from larval to embryonic stages. In particular, this work suggests cellular replacement strategies may have substantial impact in targeting the microglia, such as macrophage or haematopoietic stem cell transplantation.

## Acknowledgements

We thank Andrew Grierson for providing comments on the manuscript, David Drew, Catherine Loynes and the staff from the flow cytometry facility at the University of Sheffield, The Wolfson Light Microscopy Facility and the BSU Zebrafish for their technical help and support. Imaging work was performed at the Wolfson Light Microscopy Facility, using the Nikon W1 spinning disk confocal microscope (BBSRC grant number BB/V019368/1). The authors would also like to thank Katy Reid (formerly of the Seiger lab) for sharing her 4C4 antibody purification expertise and Ziqi Zhou for her input on the tissue clearing protocol. RNA-Seq experiment and analysis were performed with support from the Genomics Laboratory and Data Science Hub of the University of York Bioscience Technology Facility.

## Declarations

### Ethics approval

All procedures involving zebrafish followed the Animal [Scientific Procedures] Act 1986 under the Home Office Project Licence (PPL P254848FD).

## Author contributions

Conceptualization: S.A.R., N.H.; Methodology: N.H., H.A.R; Validation: H.A.R; Formal analysis: H.A.R.; Investigation: N.H., H.A.R; Resources: S.A.R., N.H.; Data curation: H.A.R., D.C.; Writing - original draft: H.A.R, N.H., S.A.R; Visualisation: H.A.R; Supervision: S.A.R., N.H., C.J.A.D.; Project administration: S.A.R., N.H.; Funding acquisition: S.A.R., N.H.

---

**Supplementary Figure 1.**
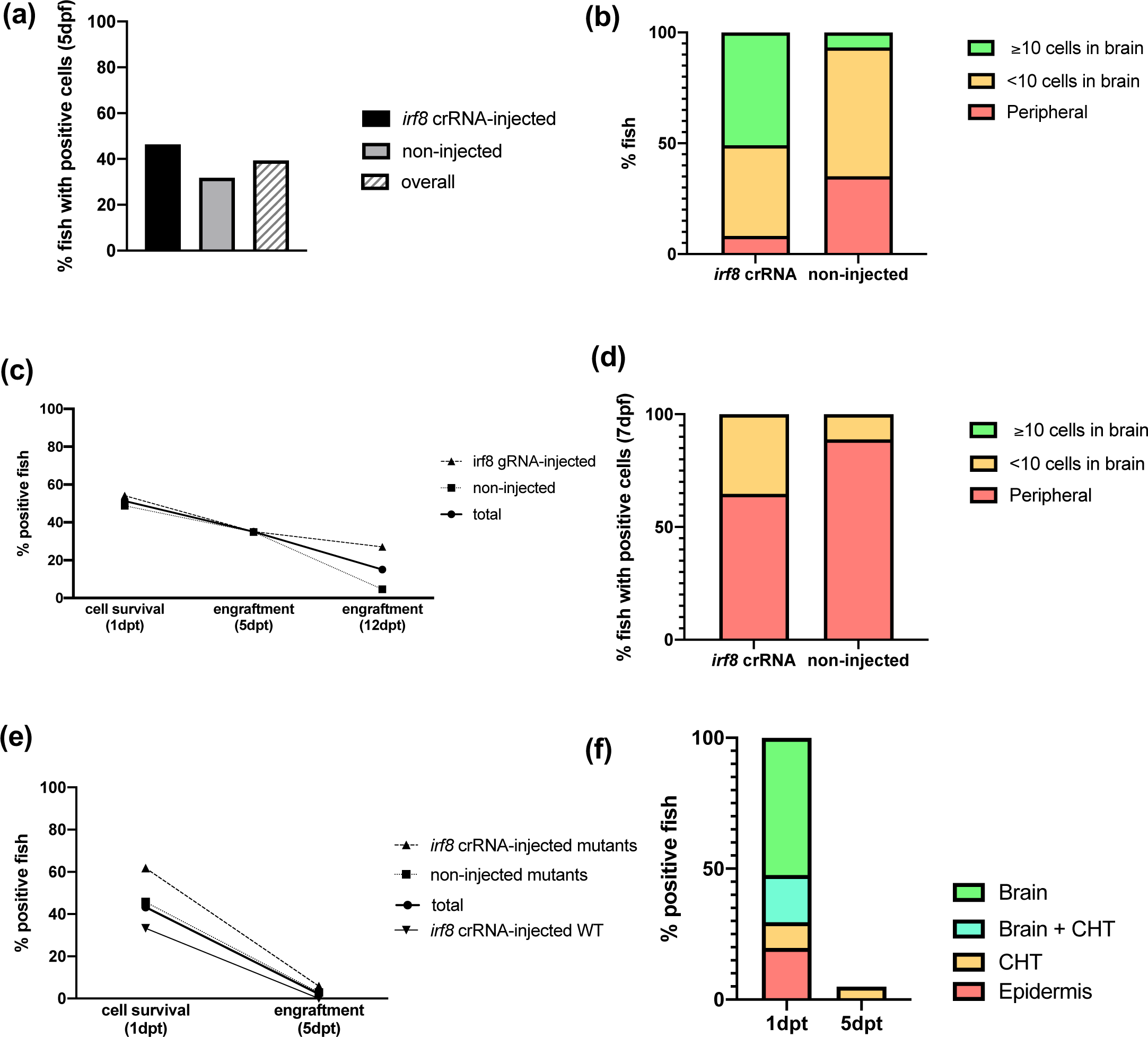
Comparison of engraftment efficiency across multiple graft sources. **a, b.** Percentage of fish with GFP-positive cells in the body **(a)** and brain **(b)** following systemic injection of *fms:GFP* cells from adult whole kidney marrow (*Tg(mpeg:mCherry CAAX)sh378* hosts, 5dpf. 3 biological replicates, n=384–423). **c, d.** Percentage of fish with GFP-positive cells in the body **(c)** and brain **(d)** following systemic injection of *CD41:GFP* cells from adult whole kidney marrow (*Tg(mpeg:mCherry CAAX)sh378* hosts, 7dpf. 1 biological replicate, n=37–43). **e, f.** Percentage of fish with GFP-positive cells in the body **(e)** and brain or CHT **(f)** following systemic injection of *fms:GFP* cells from whole embryo graft (*rnaset2^sh532^* mutant and wild type sibling hosts, 7dpf. 1 biological replicates, n=34–72). dpf, days post-fertilization. dpt, days post-transplant.

**Supplementary Figure 2.**
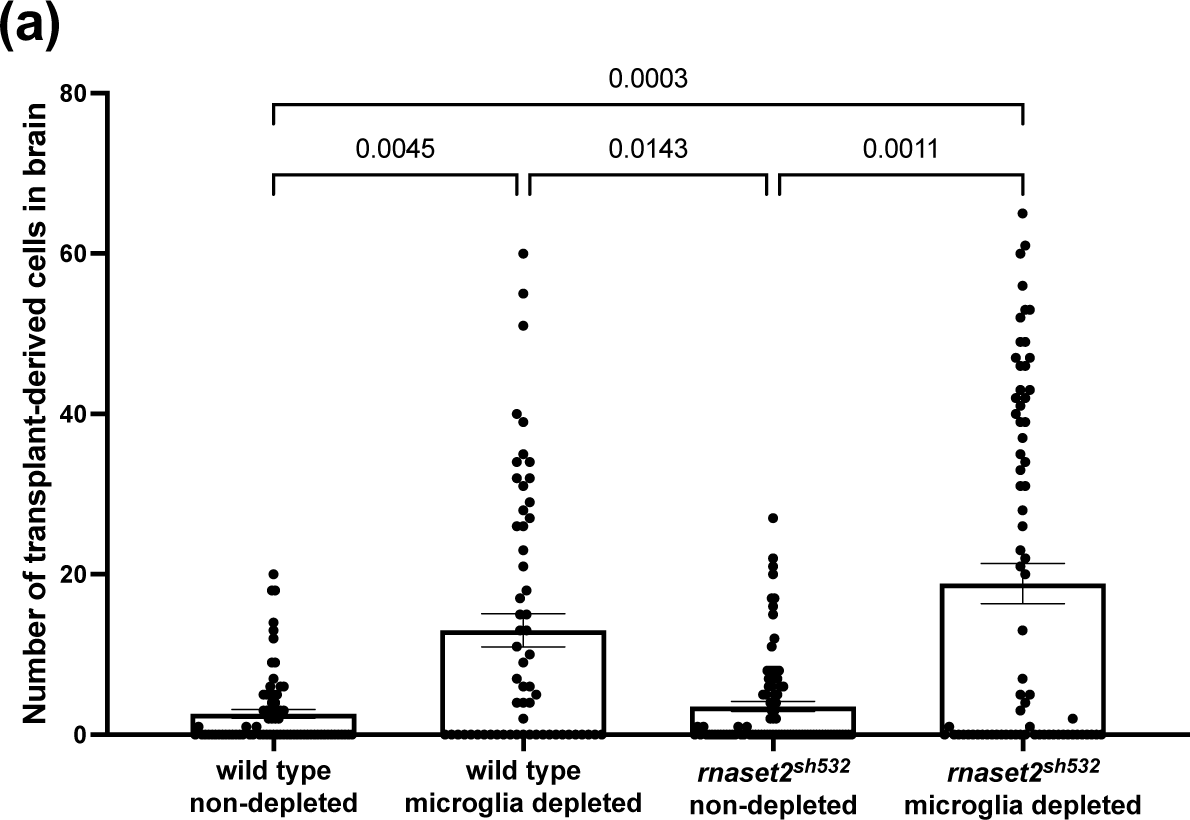
Transplanted macrophages compete with endogenous cells to fill the microglia niche in *rnaset2* mutants. **a.** Quantification of the number of GFP-positive cells within *rnaset2* mutant and wild type host brains with and without microglia-depletion via knockout of *irf8*. Kruskal-Wallis test with Dunn’s multiple comparisons, 3 biological replicates, n=60–90

**Supplementary Figure 3.**
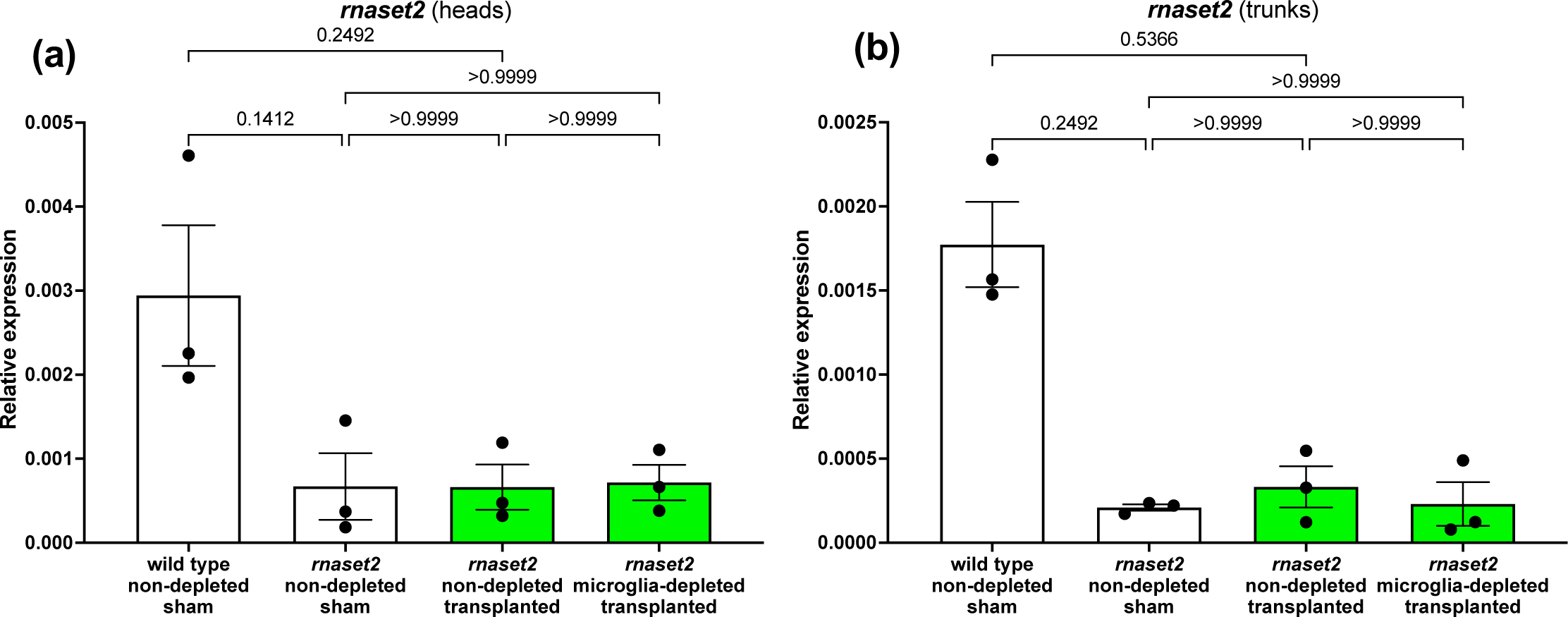
Transplantation of healthy microglia does not increase cross-correction of rnaset2 in *rnaset2* mutant head (a) nor trunks (b). Kruskal-Wallis test with Dunn’s multiple comparisons. 3 biological replicates, 15 embryos pooled per replicate.

**Supplementary Figure 4.**
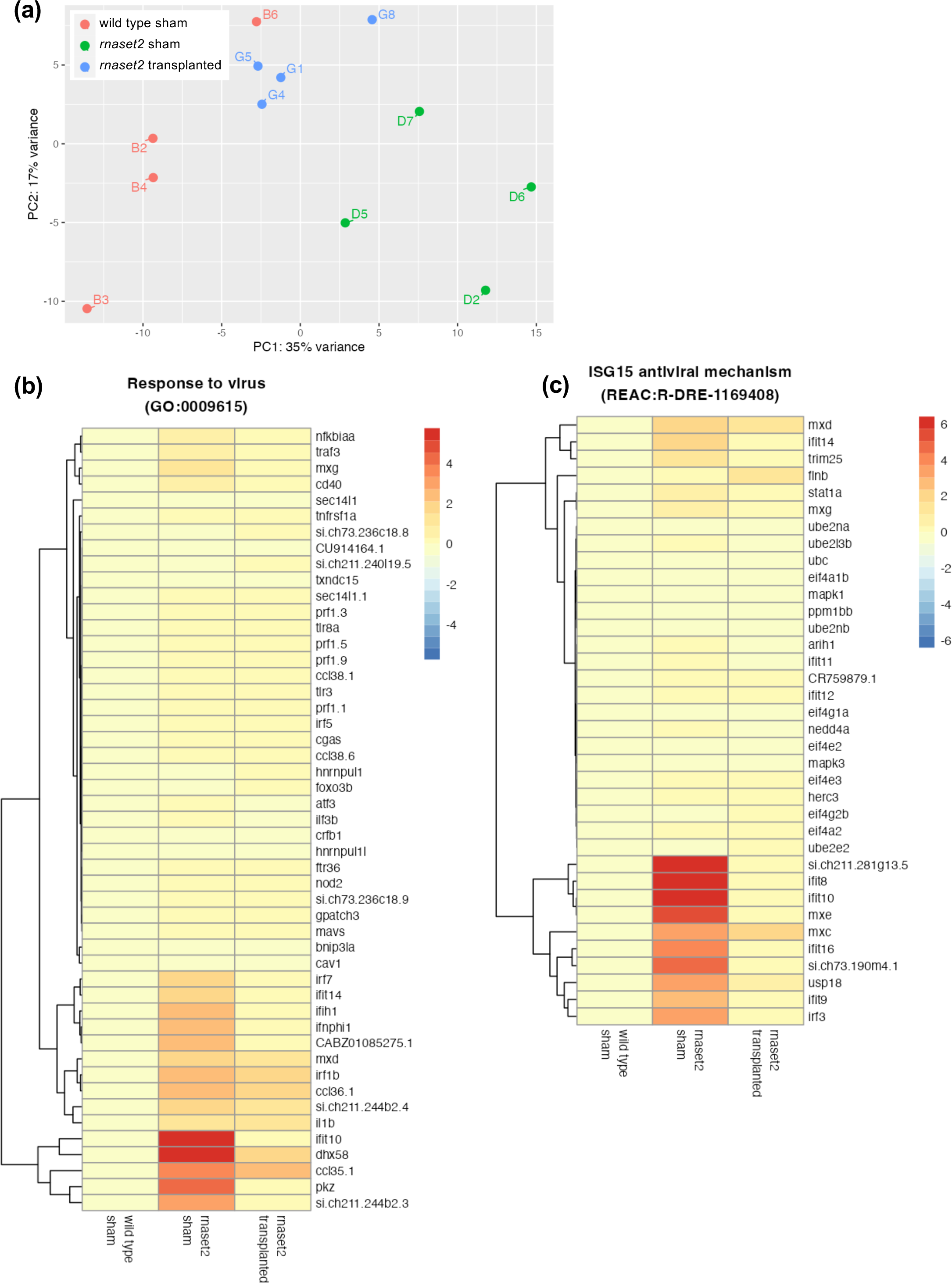
RNA sequencing of microglia-depleted wild type sham, *rnaset2^sh532^* sham and *rnaset2^sh532^* transplanted brains. **a.** Principle component analysis (PCA) plot reveals clustering of brains within each group. **b, c.** Complete heatmap of genes belonging to response to virus **(b)** and ISG15 antiviral mechanism **(c)** GO pathways for microglia-depleted animals. Fold change relative to wild type sham is indicated by color, with red indicating higher expression.

**Supplementary Figure 5.**
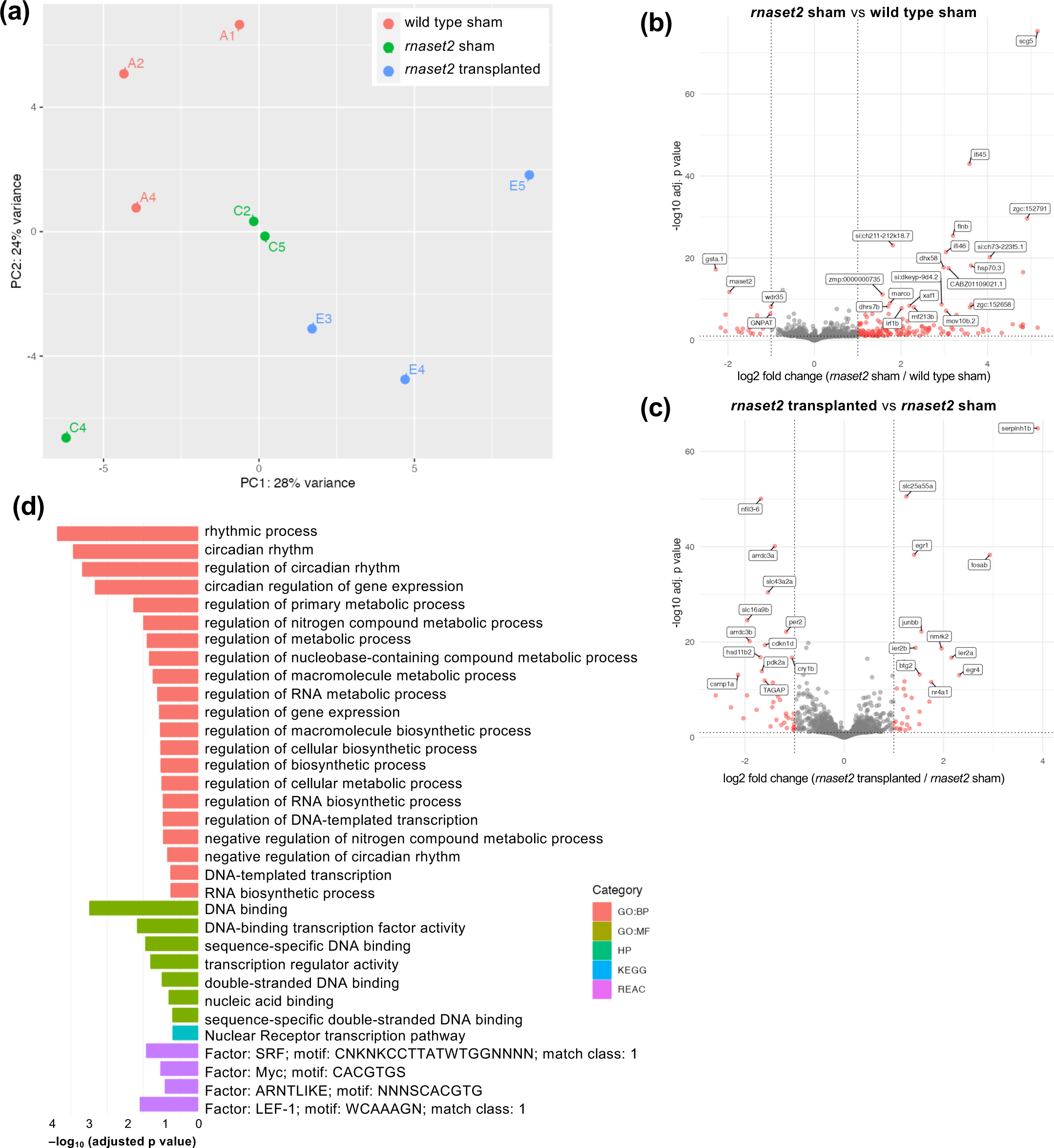
RNA sequencing of non-microglia-depleted wild type sham, *rnaset2^sh532^* sham and *rnaset2^sh532^* transplanted brains. **a.** Principle component analysis (PCA) plot reveals clustering of brains within each group. **b, c.** Volcano plot of differentially expressed genes between *rnaset2* sham versus wild type sham **(b)** and *rnaset2* transplanted versus *rnaset2* sham groups. Significantly differentially regulated genes are shown in red. The top 25 differentially expressed genes are annotated. **d.** Gene ontology (GO) plot showing significantly enriched pathways in *rnaset2* non-depleted transplanted animals compared with non-depleted sham controls, identified by gProfiler.

**Supplementary Figure 6.**
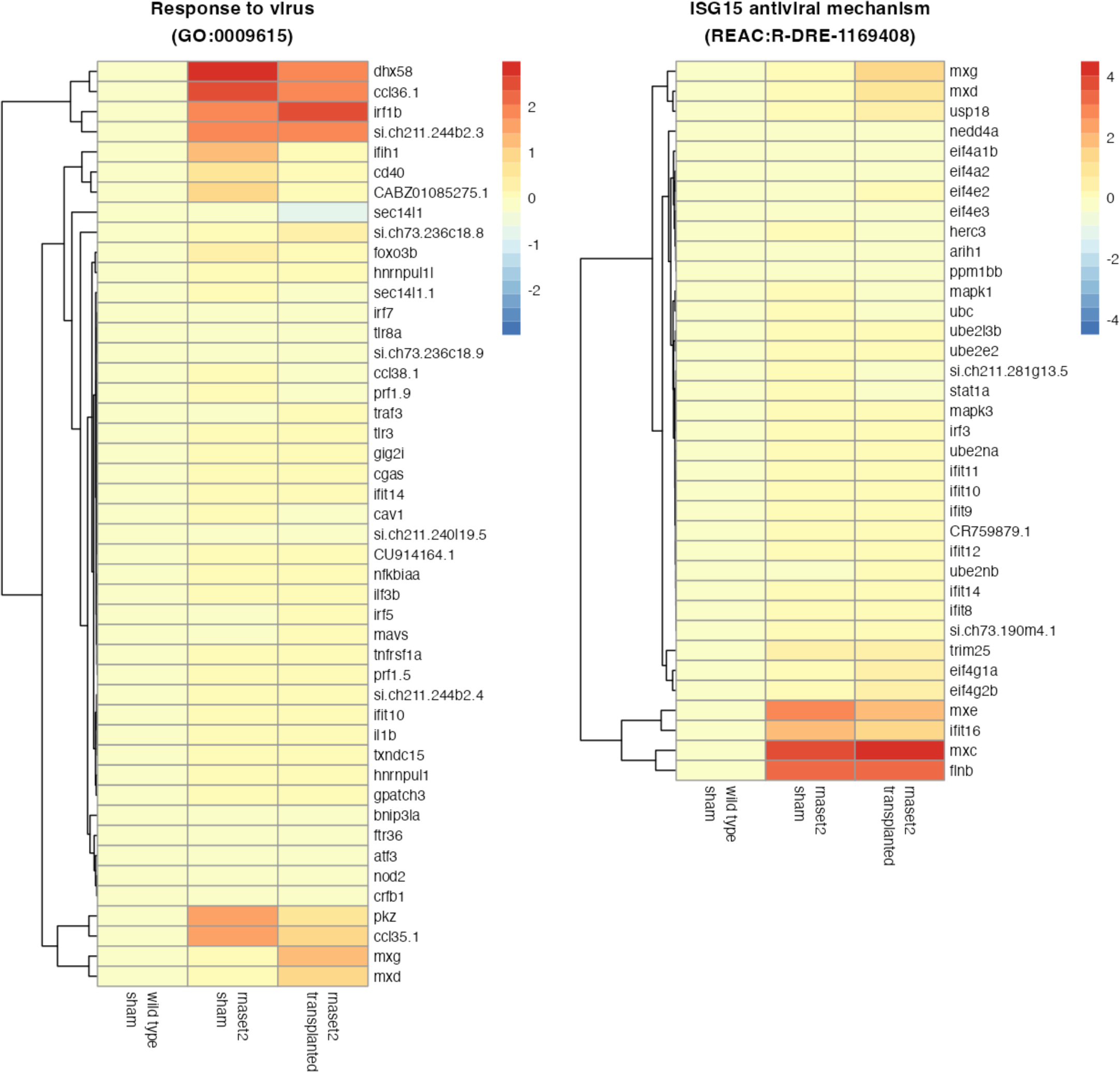
Complete heatmap of genes belonging to response to virus and ISG15 antiviral mechanism GO pathways for nondepleted animals. Fold change relative to wild type sham is indicated by color, with red indicating higher expression.

**Table S1.**
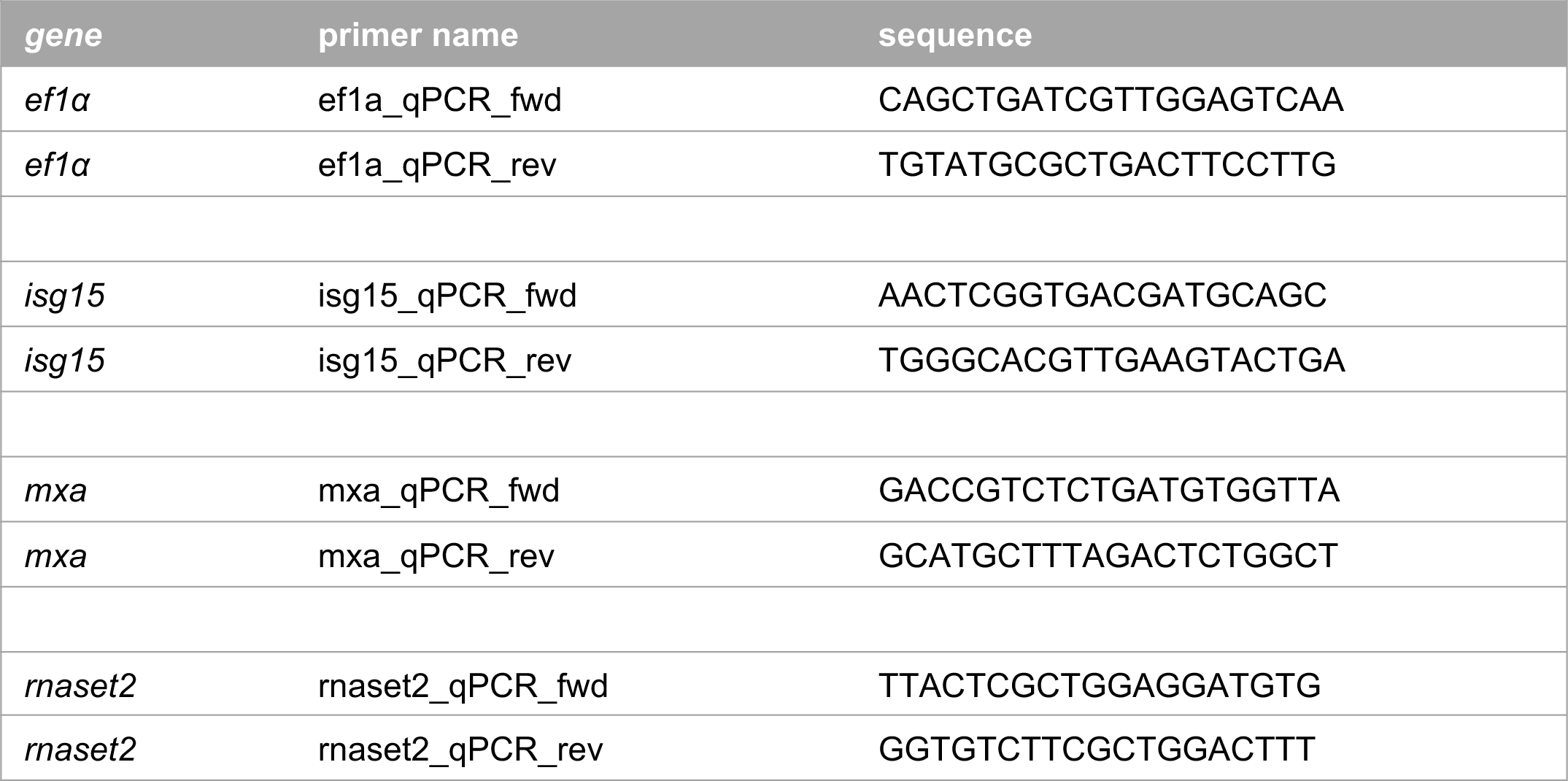
qPCR primers.

